# A mouse model of recurrent myocardial infarction reports diminished emergency hematopoiesis and cardiac inflammation

**DOI:** 10.1101/659359

**Authors:** Sebastian Cremer, Maximilian J. Schloss, Claudio Vinegoni, Shuang Zhang, David Rohde, Paolo Fumene Feruglio, Stephen Schmidt, Greg Wojtkiewicz, Ralph Weissleder, Filip K. Swirski, Matthias Nahrendorf

## Abstract

Recurrent MI is common in patients with coronary artery disease and associates with high mortality. Here we developed a surgical mouse model in which two subsequent MIs affect different left ventricular regions in the same mouse. Recurrent MI was induced by ligating the left circumflex followed by the left anterior descending branch of the coronary artery. We characterized the resulting ischemia by whole-heart fluorescent coronary angiography after optical organ clearing and by cardiac MRI. We report that a first MI induces bone marrow “memory” via a circulating signal, thereby affecting hematopoietic factor expression in bone marrow macrophages. This altered the organism’s reaction to subsequent events. Inspite at least similar extent of injury reported by blood troponin, recurrent MI caused reduced emergency hematopoiesis and less leukocytosis than a first MI. Consequently, fewer leukocytes migrated to the ischemic myocardium. The hematopoietic response to lipopolysaccharide was also mitigated after a previous MI. Our data suggest that hematopoietic and innate immune responses are shaped by a preceding MI.

## Introduction

Recurrent myocardial infarction (MI) is a common clinical problem and an independent predictor of morbidity and mortality in patients (Thune et al., 2011; Stone et al., 2014). Even with contemporary primary percutaneous intervention, reinfarction still occurs in 1 of 10 patients. One-year mortality in patients with recurrent MI is about four times higher than in patients with a first infarct (38.3% versus 10.3 %), and 20% of patients with recurrent MI die within seven days(Thune et al., 2011). Mortality is at its peak within the first month after the index MI(Thune et al., 2011). The poor outcome after recurrent MI is attributed to additional loss of viable myocardium and development of heart failure, but mechanistic studies investigating the underlying causes are lacking.

Myocardial infarction is a sterile injury that elicits an immune response to replace necrotic cardiac cells with a robust scar (Nahrendorf, 2018). A hallmark of this reaction is massive recruitment of short-lived neutrophils and monocytes from hematopoietic organs, which increase myeloid cell production by a process called emergency hematopoiesis (Nahrendorf et al., 2007; Swirski et al., 2009; Dutta et al., 2015b). Newly made leukocytes enter the heart and deploy a variety of molecular cues, namely cytokines, proteases and angiogenic substances, to enable myocardial healing. A sufficient inflammatory reaction is necessary for optimal healing after MI; however, unrestrained myeloid cell activity results in adverse cardiac remodeling and heart failure (Frangogiannis, 2012).

Recently the concept of “trained immunity” (Netea et al., 2016) reinvigorated interest in changes to innate immune cells’ functional programs. After inflammatory stimuli, i.e. “training”, more vigorous activity occurs in response to subsequent events, while states of tolerance occur after lipopolysaccharide challenge (Foster et al., 2007; Saeed et al., 2014). The initial findings obtained in macrophages have now been extended to bone marrow hematopoiesis and “training” with IL-1β and atherogenic diet (Christ et al., 2018; Mitroulis et al., 2018). Conceptually, this body of work indicates that the innate immune system may react non-uniformly depending on prior challenges to the equilibrium. Within this framework, the innate immune response to MI may also depend on prior stimuli, yet it is currently unclear whether a second infarct elicits a response identical to that observed after a first MI. Motivated by the importance of leukocyte recruitment after MI for acceleration of atherosclerosis and for heart failure development (Nahrendorf, 2018), we hypothesized that the inflammatory signals inducing emergency hematopoiesis after MI also have consequences for recurrent myocardial ischemia.

Since there was no method available to test this hypothesis, we developed a model of recurrent MI, which we describe for the first time in this Resource Article. In the mouse, the major part of the left ventricle is supplied by the left coronary artery, which emerges under the left atrium and continues to the apex of the heart (Fernández et al., 2008). It begins as a main trunk of 2-3mm length. Below the tip of the left atrial appendage, the artery divides into i) a branch with significant caliber that supplies blood to the posterolateral area of the heart (left circumflex, LCX)(Ahn et al., 2004; Fernández et al., 2008); ii) a branch that continues to the apex of the heart and provides blood to the anterior wall of the heart (left anterior descending, LAD)(Ahn et al., 2004; Fernández et al., 2008). We first ligated the LCX and later separately ligated the LAD branch to cause a second MI in the same mouse. We used whole heart fluorescence coronary angiography (FCA) and delayed gadolinium enhancement magnetic resonance imaging (MRI) to map the sequelae of recurrent MI. We then tested the hypothesis that recurrent MI induces an immune response that diverges from a first infarct.

## Results

### A mouse model of recurrent MI

Investigating recurrent multifocal ischemia in mice requires a method that allows subsequent MIs at different locations in the same mouse heart. The left coronary artery’s first branching point gives rise to the left anterior descending artery (LAD) and a circumflex artery (LCX) (Fernández et al., 2008). Since both arterial branches are fairly large caliber, we devised a model in which each is separately ligated distal of the branching point (**Figure 1A, B**). To explore both surgeries’ outcome, we performed whole heart fluorescent coronary angiography (FCA) after optical tissue clearing (Richardson and Lichtman, 2015) and in vivo labeling of the cardiac vasculature with a fluorescent dye. Without tissue clearing, coronary arteries were not visible because they are partially covered by myocardial tissue, which efficiently absorbs and scatters photons (data not shown). We therefore adapted CUBIC (clear, unobstructed brain imaging cocktails and computational analysis), a tissue clearing method previously developed for brain imaging (Susaki et al., 2014). Tissue clearing was done after injection of fluorescein-labelled albumin, which served as a fluorescent vascular imaging agent (see method section). FCA clearly visualized the coronary arteries and their branches LAD and LCX (**Figure 1C**). Ligating the LCX branch induced a strictly posterolateral infarct that did not extend to the apex of the heart (**Figure 1D and Figure S1**), while the LAD ligation distal of the main branching point affected a large portion of the anterior wall including the apex **(Figure 1E**). Finally, after both ligations, a very large infarct covering a major area of the left ventricle emerged **(Figure 1F, Figure S1).**

**Figure 1.**
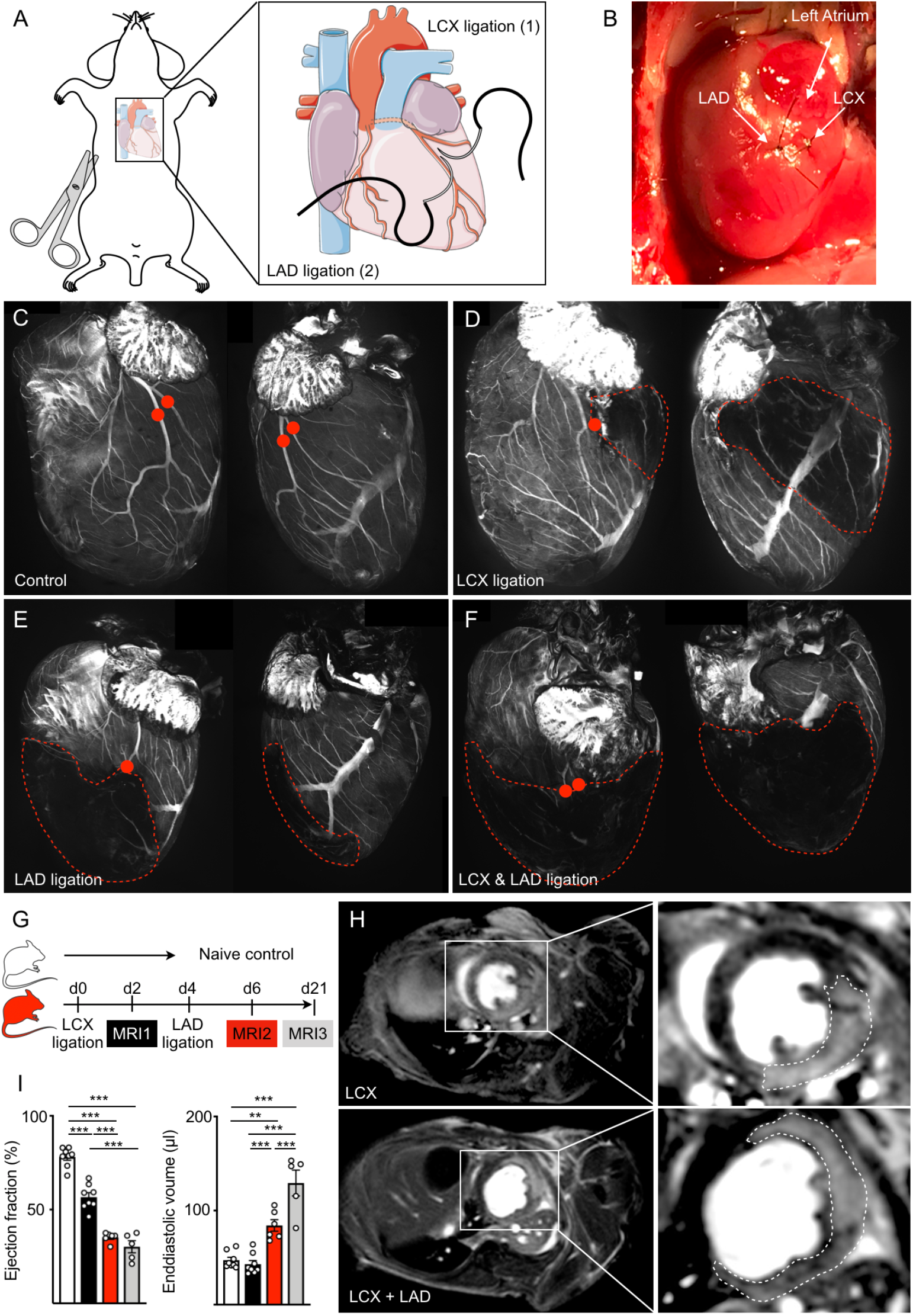
Recurrent MI in mice. (**A**) Surgical approach. The LAD and its first branch (LCX) are ligated separately. (**B**) Image of LAD and LCX ligations in the same heart. (**C**) Confocal projections of the coronary vasculature after optical clearing and in vivo staining with fluorescein albumin in naive mice, (**D)** mice with LCX and **(E)** LAD infarctions and **(F)** after ligation of both vessels. Red dots indicate the ligation sites; the infarcted area is outlined. **(G)** Timeline of MRI. (**H**) MRI delayed enhancement short axis views of a mouse with LCX MI (upper panel) and the same mouse after subsequent LAD MI (lower panel). Insets show magnification views of the left ventricle. The MI is indicated by a dotted line. (**I**) Ejection fraction and end-diastolic volume, assessed by MRI, of naive mice and mice after one or two MIs and three weeks thereafter. **p<0.01, ***p<0.001, n=5-8, one-way analysis of variance (ANOVA) followed by Tukey’s multiple comparisons test. Data are mean ± s.e.m..

To examine the consequences of these surgeries, we conducted a cardiac MRI study with delayed gadolinium enhancement, a gold standard for identifying non-viable myocardium in mice and humans (Kim et al., 2000; Yang et al., 2004) (for timeline, see **Figure 1G**). LCX ligation created posterolateral, gadolinium-enhanced infarcts that significantly reduced the ejection fraction two days after MI, while the dimensions of the left ventricle remained unchanged at this early time point after ischemia **(Figure 1H, I and supplementary video 1).** Additionally infarcting the LAD in the same mouse two days later generated larger infarcts, reduced the left ventricular ejection fraction and induced dilation of the left ventricle **(Figure 1H, I and supplementary video 2)**. Twenty-one days after the first MI (17 days after the second), the left ventricle dilated further (**Figure 1I**).

### Hematopoietic response to a single LAD versus a single LCX infarct

Appropriate leukocyte supply is necessary to promote optimal healing of the infarcted heart, as recruited immune cells remove dead myocytes and orchestrate wound healing (Swirski et al., 2009). Most recruited cells are innate immune cells, with short half lives, that are supplied by hematopoietic tissues. MI activates bone marrow to generate a sufficient leukocyte supply to the heart in mice and humans(Dutta et al., 2015b). We first tested whether lateral infarcts after LCX ligation induce a bone marrow activation similarly to previously described LAD infarcts (Dutta et al., 2015b). Comparing lineage^−^ ckit^+^ sca-1^+^ (LSK) progenitor cells and most upstream CD150^+^ CD48^+^ hematopoietic stem cell (SLAM HSC) numbers in the mice femurs after LCX ligation versus LAD ligation or naive controls, we found that HSPC numbers increased after LCX ligation when compared to naive controls, albeit not to the same extent as after LAD ligation **(Figure 2A, B)**, likely because LCX infarcts are smaller than LAD infarcts. The number of downstream granulocyte macrophage progenitors (GMP) increased robustly after either LCX or LAD ligation (**Fig. 2B**).

**Figure 2.**
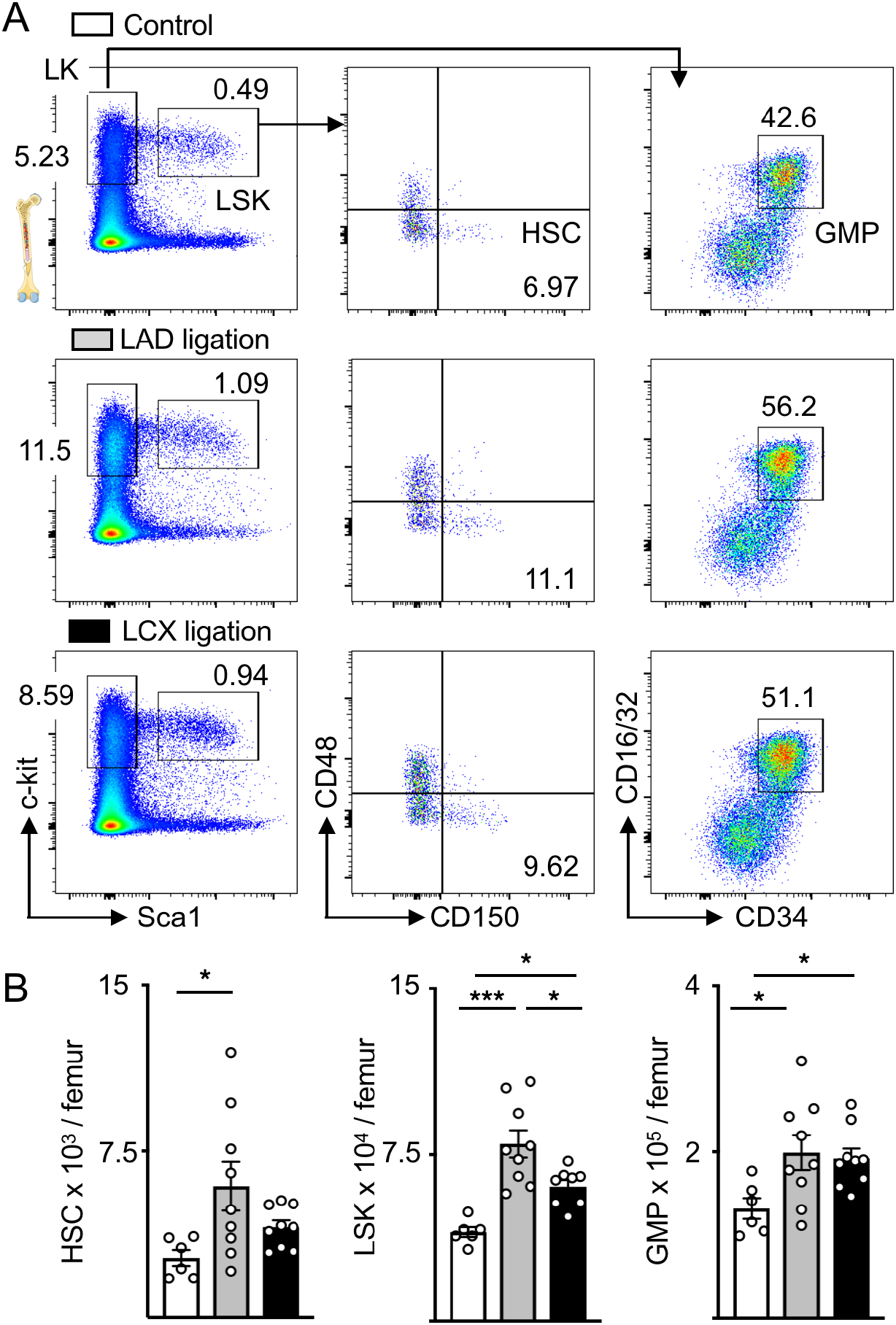
LCX and LAD ligation activate the bone marrow. **(A)** Dot plots of hematopoietic stem and progenitor cells (LT-HSC, LSK and GMP) in naive mice, mice three days after LAD or LCX ligation. **(B**) Quantification of cell numbers of LT-HSC, LSK and GMP. *p<0.05, ***p<0.001, n=6-9, one-way analysis of variance (ANOVA) followed by Tukey’s multiple comparisons test. Data are mean ± s.e.m..

### Recurrent MI triggers attenuated systemic inflammation

It was unclear whether prior MI alters the hematopoietic emergency response to a second infarct. To investigate, we designed an experiment that compares the inflammatory response after permanent LAD ligation in mice that either had or had not received a LCX ligation 10 days before **(Figure 3A).** We first evaluated whether emergency hematopoiesis had returned to steady state 10 days after MI **(Figure S2A)**. Indeed, at this time point, the bone marrow’s hematopoietic stem and progenitor cell (HSPC) numbers, which expand significantly in the days after MI (Dutta et al., 2015b), had dropped to normal levels **(Figure S2B, C).** Likewise, neutrophil and monocyte numbers were comparable in naive mice and mice 10 days after LAD ligation **(Figure S2D)**. We concluded that a 10-day interval between the LCX and LAD ligation is appropriate because i) at that point hematopoietic cell numbers have returned to steady state, ii) the time interval is clinically relevant (Thune et al., 2011; Stone et al., 2014) and iii) a comparable time interval has previously been used to study “trained immunity” (Saeed et al., 2014; Mitroulis et al., 2018). We thus settled on an experimental design in which a control cohort receives only LAD ligation, while the experimental cohort receives an LAD ligation 10 days after a preceding LCX infarct (recurrent MI, **Figure 3A**). Blood troponin levels indicated that the acute myocardial damage caused by permanent LAD ligation was not smaller in mice with prior LCX ligation **(Figure S3).**

**Figure 3.**
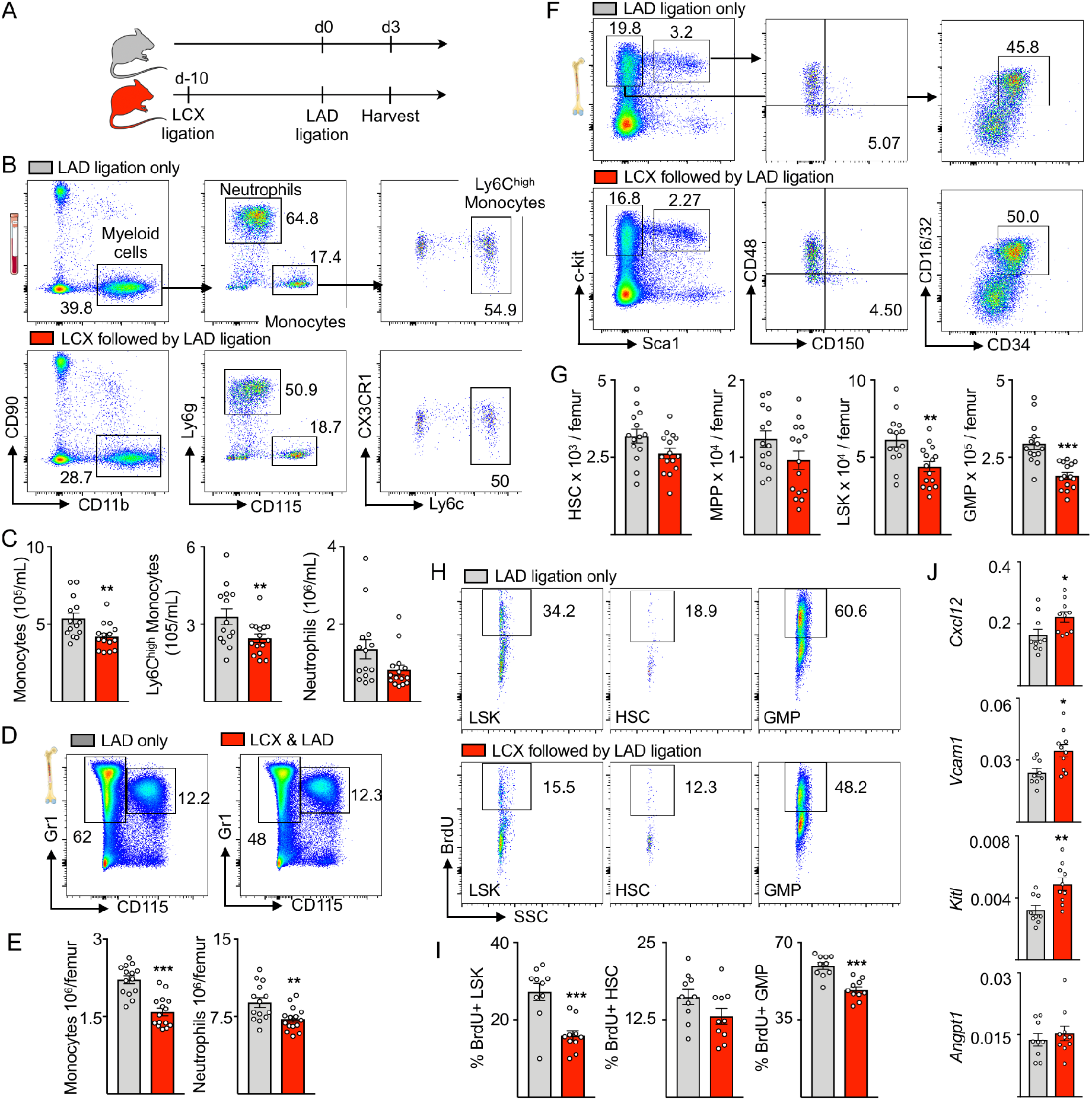
Attenuated systemic inflammation in recurrent MI. **(A)** Experimental outline. Mice were subjected to LCX MI 10 days before LAD MI. Mice were analyzed on day three after LAD MI. **(B)** Dot plots and **(C)** quantification of blood myeloid cells from mice with only one MI compared to recurrent MI. **(D)** Dot plots and **(E)** quantification of neutrophils and monocytes in bone marrow. **(F)** Flow plots and **(G)** quantification of HSPC in bone marrow. **(H)** Dot plots and **(I)** quantification of BrdU incorporation for HSPC proliferation. **(J)** Gene expression by qPCR in total bone marrow. mRNA levels were normalized to *gapdh* Ct values. *p<0.05, **p<0.01, ***p<0.001, n=9-15, Student’s t-test. Data are mean ± s.e.m..

We next compared innate immune cell numbers 3 days after a first MI (LAD only) to 3 days after recurrent MI (LAD with preceding LCX). Surprisingly, blood monocyte and neutrophil numbers were lower in mice with recurrent MI (**Figure 3B, C).** This was accompanied by lower levels of neutrophils and monocytes in the bone marrow **(Figure 3D, E)**, indicating leukocyte production — rather than only release — is reduced in the marrow. Indeed, LSK and GMP numbers were significantly lower on day 3 after recurrent MI **(Figure 3F, G)**, when post-MI emergency hematopoiesis peaks (Dutta et al., 2015b). In alignment with these HSPC numbers, LSK and GMP proliferation was lower in the femurs of mice with recurrent MI than after a first infarct, as shown by analyzing cell proliferation using a BrdU incorporation assay **(Figure 3H, I**). This trend did not reach statistical significance for upstream SLAM HSC **(Figure 3H, I**).

### Recurrent MI changes the hematopoietic niche

In addition to growth factors and danger signals, HSPC proliferation and leukocyte production is governed by the hematopoietic niche, specifically by proteins encoded by the genes *cxcl12, angpt1, kitl* and *vcam1* (Mendelson and Frenette, 2014; Morrison and Scadden, 2014). Expression of these genes was significantly higher in mice with recurrent MI compared to those with a first infarct, except for *angpt1* **(Figure 3J).** These data implicate the involvement of the bone marrow niche, which instructs HSPC activity, in our observation of lower hematopoietic expansion after recurrent MI.

### Preceding ischemia reperfusion injury restricts post-MI emergency hematopoiesis

Next we extended the above findings to ischemia/reperfusion injury (IRI). We instigated ischemia by blocking LAD branch blood supply for 30 minutes, followed by reperfusion of the occluded vessel. Ten days later mice underwent permanent ligation of the LAD while controls received LAD ligation without preceding IRI **(Figure 4A)**. Troponin levels in mice in which IRI preceded MI were not below those of control mice that only underwent LAD ligation **(Figure 4B).** Similar to the observations in recurrent MI described above, mice with prior IRI had lower bone marrow LSK and GMP numbers three days after permanent LAD ligation **(Figure 4C, D).** Likewise, HSPC proliferation was significantly lower as assessed by BrdU incorporation **(Figure 4E, F).** These data indicate that a “memory” of the bone marrow compartment can be induced not only by non-reperfused but also reperfused MI.

**Figure 4.**
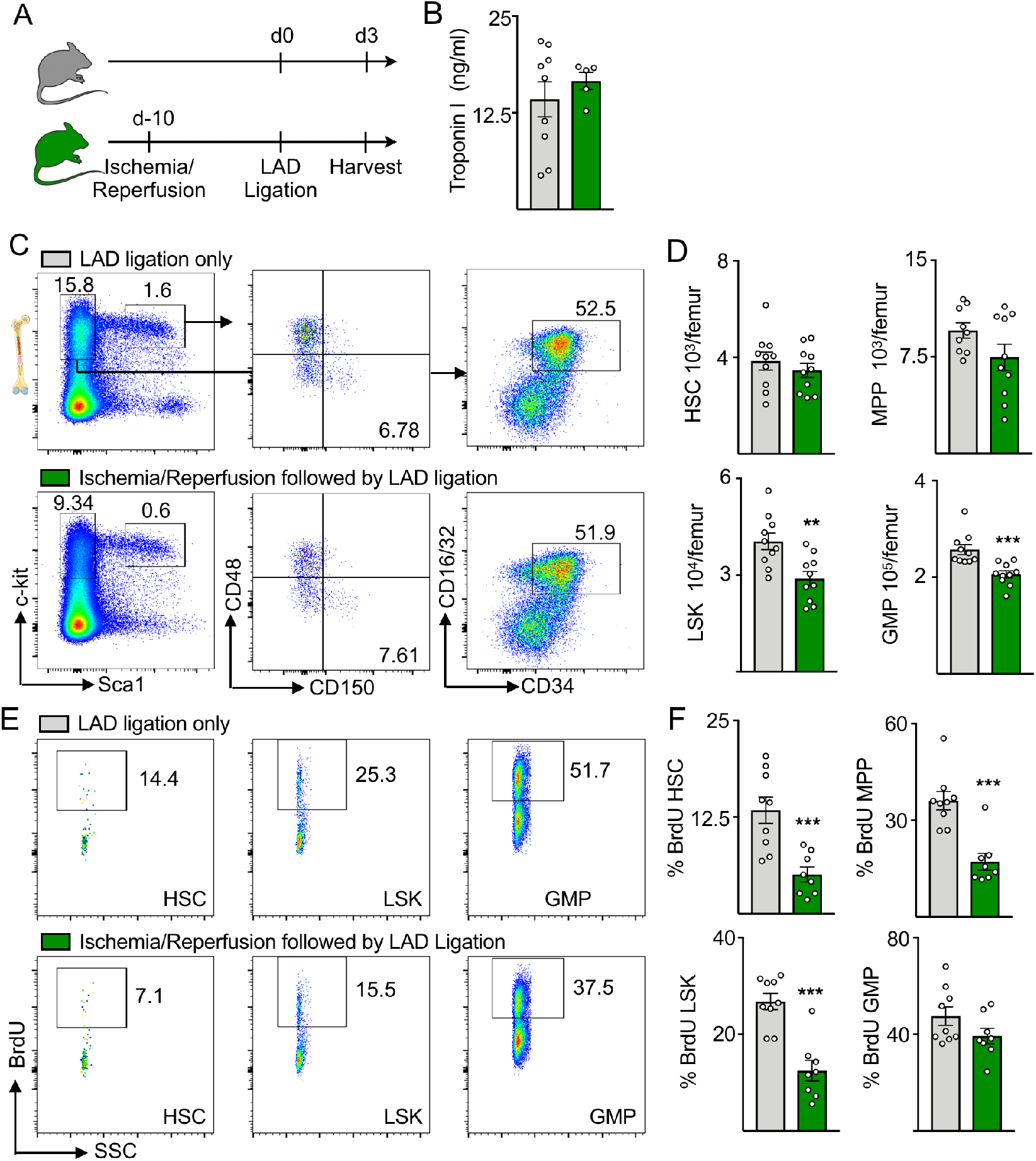
Previous ischemia reperfusion injury (IRI) restricts bone marrow activity after MI. **(A)** Experimental design. Mice were subjected to 30 min ischemia by temporary LAD ligation followed by reperfusion 10 days before inducing LAD MI. Mice were analyzed on day three after LAD MI. **(B)** Troponin-I values 24 hours after LAD MI (n=4-9). **(C)** Flow plots and **(D)** quantification of HSPC numbers in bone marrow. **(E)** Dot plots and **(F)** quantification of BrdU incorporation assay for bone marrow HSPC. **p<0.01, ***p<0.001, n=8-10, Student’s t-test. Data are mean ± s.e.m..

### A circulating factor dampens emergency hematopoiesis in recurrent MI

After myocardial ischemia, the bone marrow is activated by various mechanisms, such as direct signaling of sympathetic nerves (Dutta et al., 2012) or by soluble factors, which are produced in the injured myocardium (Sager et al., 2015). To test whether a circulating factor limits emergency hematopoiesis after recurrent MI, mice were joined in parabiosis to create a shared circulation. After two weeks of parabiosis, the time necessary to create equilibrium between both mice, one parabiont received an LAD MI while the other parabiont did not. Five days later, mice were split and the non-infarcted parabiont was subsequently subjected to an LAD MI in both cohorts. Mice in the experimental group were therefore exposed to the blood of the infarcted parabiont 10 days prior to MI. Mice joined in parabiosis without prior MI were given LAD ligation and used as controls **(Figure 5A).**

**Figure 5.**
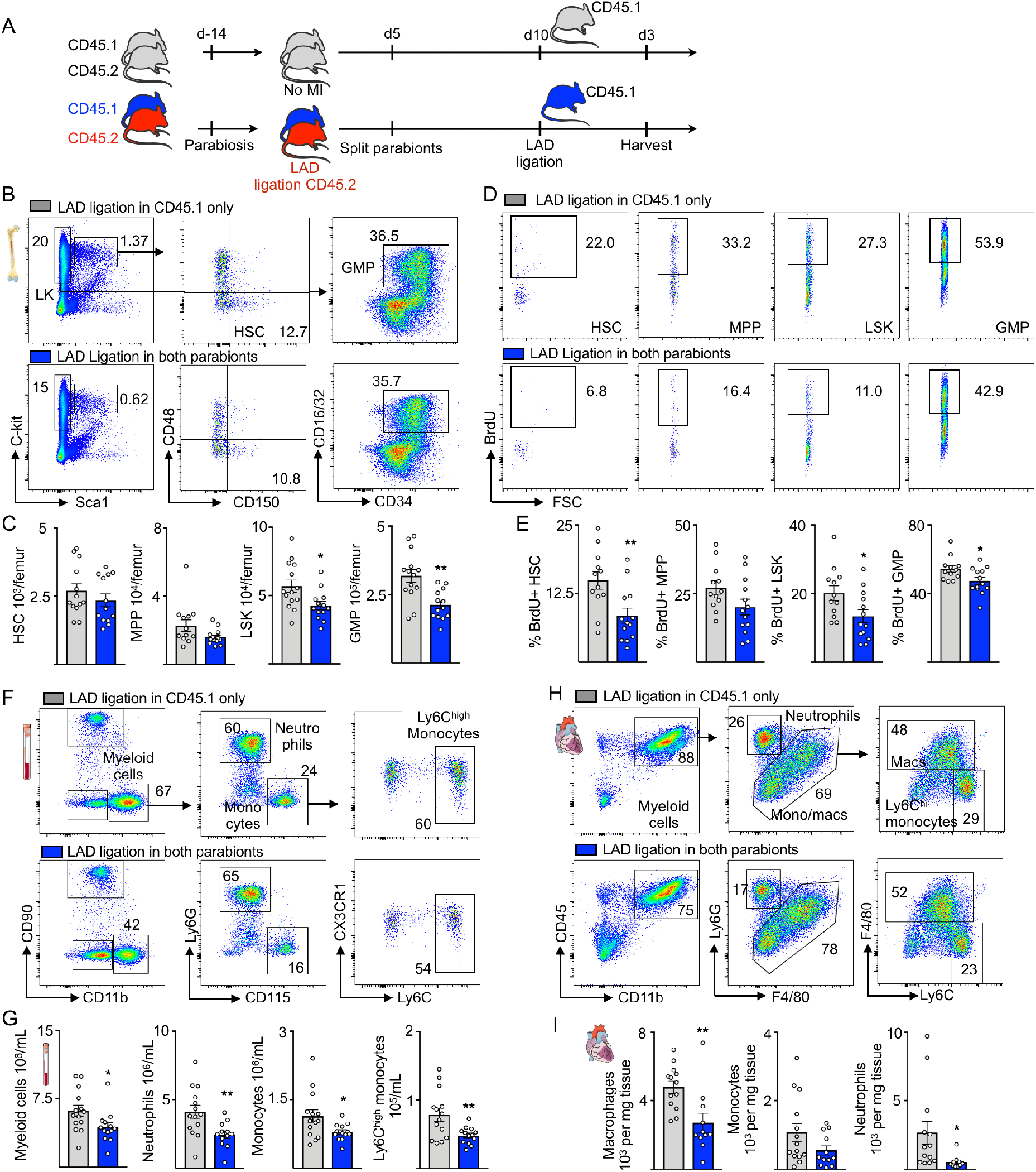
A circulating factor attenuates progenitor cell expansion. **(A)** Experimental design. Mice were joined in parabiosis. Two weeks later one mouse received MI surgery. Five days later parabionts were separated, and LAD ligation was conducted in the other mouse five days after. **(B)** Dot plots and **(C)** quantification of bone marrow HSPC. **(D)** Dot plots and **(E)** quantification of BrdU incorporation assay for bone marrow HSPC. **(F)** Dot plots and **(G)** quantification of blood myeloid cells. **(H)** Flow cytometry gating and **(I)** quantification of leukocyte subsets from infarcted myocardium. *p<0.05, **p<0.01, n=12-14, Student’s t-test. Data are mean ± s.e.m..

We found that LSK and GMP numbers were lower in the bone marrow of mice in which a parabiont had previous MI surgery **(Figure 5B, C).** Accordingly, we found lower proliferation rates of LT-HSC, MPP, LSK and GMP in the bone marrow of these mice **(Figure 5D, E).** This lower proliferation resulted in decreased numbers of myeloid cells in the blood **(Figure 5F, G)**, which serves as a conduit for cells migrating to the infarct. Indeed, we detected fewer myeloid cells in the infarcted myocardium three days after MI in mice in which the parabiont had experienced prior MI **(Figure 5H, I).** Taken together, these data demonstrate that a soluble signal travels from the infarcted parabiont to the non-infarcted parabiont’s bone marrow, where it dampens bone marrow activation after a recurrent ischemic event. Hypothetically, IL-1β may have such a role, given that it alerts the marrow after MI (Sager et al., 2015) and that it has lasting effects on hematopoiesis in hyperlipidemia (Christ et al., 2018).

### Prior MI attenuates LPS-induced emergency hematopoiesis

To test if these observations extend to non-ischemic recall challenges, we conducted experiments in which the bacterial wall component lipopolysaccharide (LPS) served as a secondary challenge ten days after MI **(Figure 6A)**. This experimental design follows classical “trained immunity” experiments, which likewise rely on LPS recall (Saeed et al., 2014; Mitroulis et al., 2018). The bone marrow was analyzed 24 hours after injection of LPS, which induces severe systemic inflammation and emergency myelopoiesis to meet the organism’s demand for myeloid cells to fight bacteria (Mitroulis et al., 2018). After LPS injection, we found fewer HSC and LSK as well as significantly reduced progenitor cell numbers in the bone marrow of mice with prior MI **(Figure 6B-D).** In addition, we observed a non-significant trend towards reduced neutrophil and monocyte numbers in the bone marrow of mice that underwent MI 10 days prior to LPS exposure **(Figure 6E).** As after recurrent MI, leukocyte production was diminished, indicated by lower BrdU incorporation in HSPC of mice with MI prior to LPS injection **(Figure 6F, G).** In sum, these data indicate that ischemic injury of the heart induces “tolerance” or exhaustion in the bone marrow, preventing robust HSPC expansion and generation of sufficient amounts of innate immune cells after re-exposure to inflammatory stimuli.

**Figure 6.**
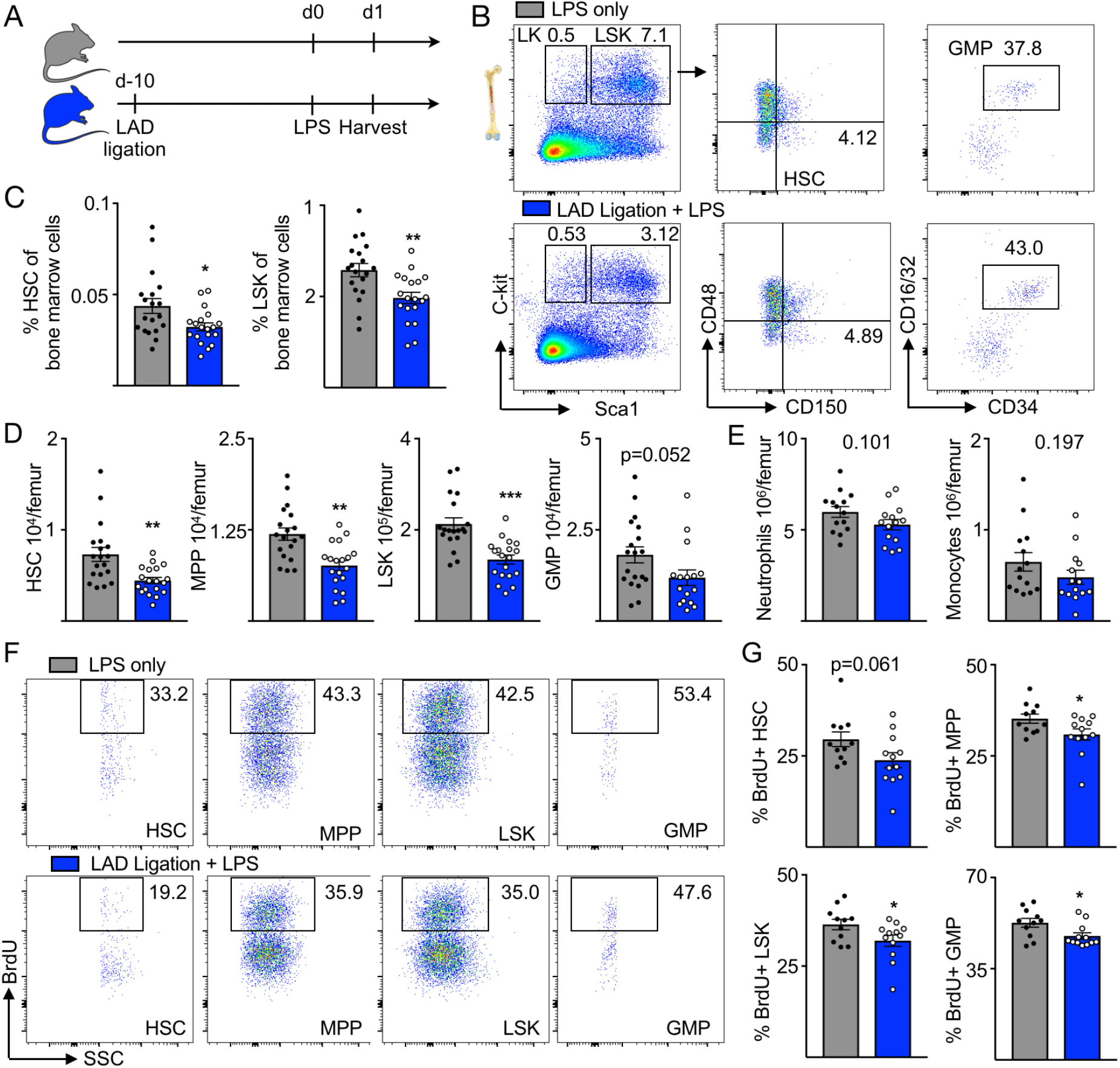
Prior MI attenuates HSPC expansion after LPS. **(A)** Experimental design. Mice received MI surgery 10 days before sepsis induction by i.p. LPS injection. **(B)** Flow plots, **(C)** frequencies and **(D)** quantification of bone marrow HSPC. **(E)** Leukocytes in bone marrow. **(F)** Representative dot plots and **(G)** quantification of BrdU incorporation for bone marrow HSPC. *p<0.05,**p<0.01, ***p<0.001, n=12-14, Student’s t-test. Data are mean ± s.e.m..

### Macrophage ablation abolishes bone marrow “tolerance”

Macrophages reside in the bone marrow and are a long-lived component of the hematopoietic niche (Chow et al., 2011; Chow et al., 2013), which governs HSPC quiescence, maintenance and proliferation. In the bone marrow, macrophages were characterized as CD3^−^ B220^−^ Gr1^−^CD115^−^ F4/80^+^ SSC^lo^. As previously described (Chow et al., 2011), we found macrophages to be MHCII^intermed^ Cx3cr1^neg^ CD169^high^ Vcam1^high^ (**Figure S4).** Given prior reports of trained immunity and tolerance (Foster et al., 2007; Biswas and Lopez-Collazo, 2009; Cheng et al., 2014; Saeed et al., 2014), we hypothesized that bone marrow macrophages may be involved in retaining “memory” of MI. To test this hypothesis, we depleted macrophages with clodronate liposomes in mice with or without prior MI **(Figure 7A)** and examined the effect of LPS on HSPC expansion. Clodronate efficiently depleted macrophages in the bone marrow **(Figure 7B, C).** We reproduced the attenuated myeloid cell expansion described above in mice with prior MI that received LPS and control liposomes compared to mice without MI (**Figure 7D, E**). After macrophage ablation, HSPC numbers fell even in the absence of MI (**Figure 7D, E**), which is in accordance with prior reports suggesting that niche macrophages maintain HSPC, for instance via the adhesion molecule Vcam-1 (Chow et al., 2011; Chow et al., 2013; Dutta et al., 2015a). Importantly, in the absence of macrophages, progenitor cell numbers after LPS challenge were similar irrespective of whether or not mice had received an MI 10 days previous. Viewed together with data on macrophages’ roles in regulating hematopoiesis in the bone marrow stem cell niche (Chow et al., 2011; Chow et al., 2013; Hashimoto et al., 2013), these results indicate that after a prior stimulus, bone marrow macrophages may indeed influence the dampened emergency hematopoiesis after recurrent MI.

**Figure 7.**
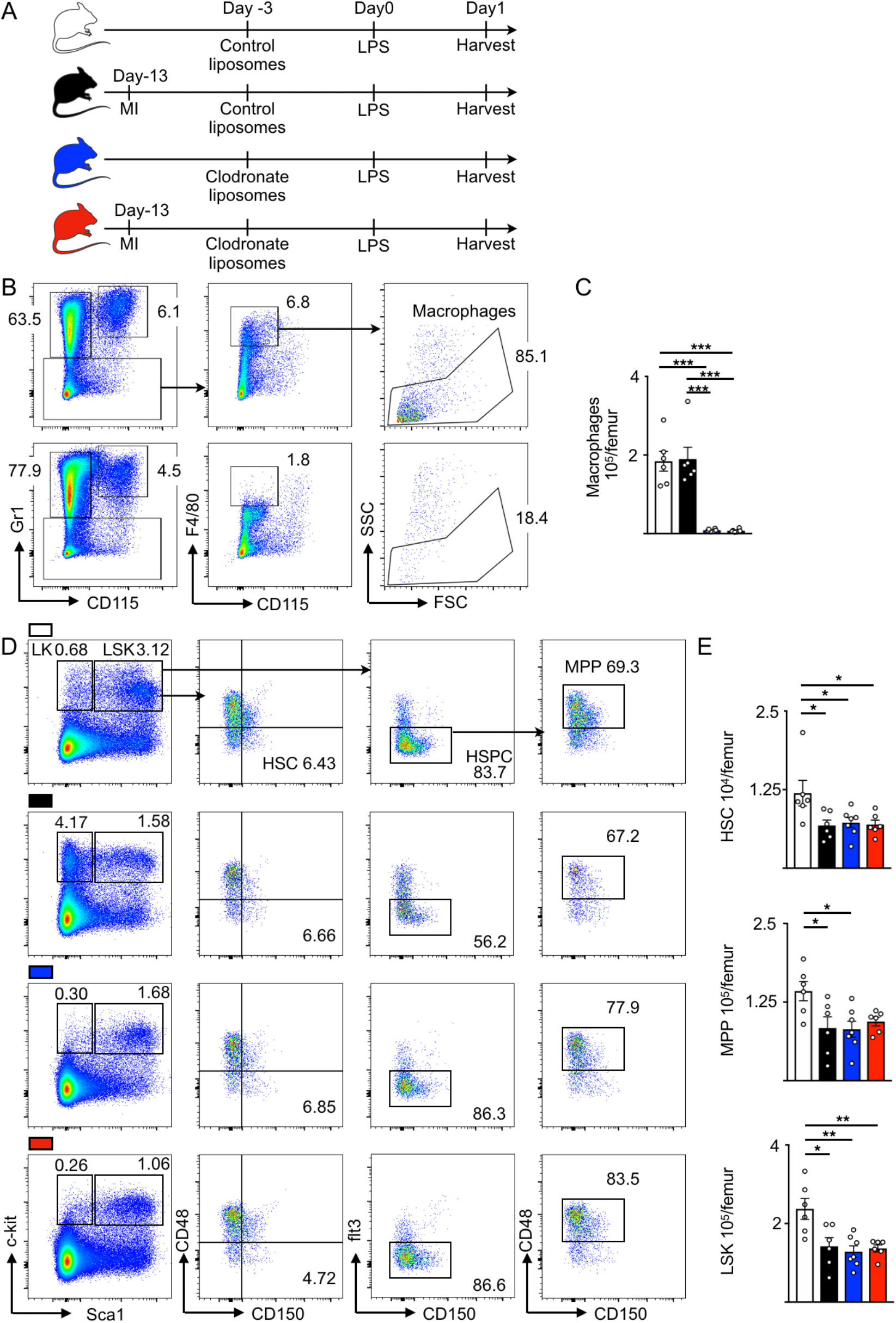
Macrophage ablation abolishes bone marrow memory. **(A)** Experimental design. Mice received either control liposomes or clodronate liposomes. Half of the mice in each cohort underwent LAD MI. All mice were injected with LPS. **(B)** Dot plots and **(C)** quantification of bone marrow macrophages showing efficient depletion by clodronate liposomes. **(D)** FACS-plots and **(E)** cell numbers of bone marrow HSPC after clodronate depletion. *p<0.05, **p<0.01, ***p<0.001, n=5-8, one-way analysis of variance (ANOVA) followed by Tukey’s multiple comparisons test. Data are mean ± s.e.m..

To analyze the changes in the bone marrow niche in more detail, we isolated key stromal hematopoietic niche cells with FACS sorting in mice three days after MI and compared niche factor expression with naive controls and mice with recurrent MI **(Figure 8A).** The flow cytometric gating for endothelial and leptin receptor^+^ (LepR-YFP^+^) stromal cell isolation is shown in **Figure 8B.** There were no significant differences for *kitl, Vcam1*, *Angpt1* and *Cxcl12* expression in LepR^+^ stromal mesenchymal cells or bone marrow endothelial cells in mice with single versus recurrent MI **(Figure 8C, D)**. However, in mice with recurrent MI, bone marrow macrophages expressed significantly more *Vcam1* and *Angpt1,* at levels comparable to naive controls **(Figure 8E)**. Since these factors regulate HSPC retention and quiescence, the data suggest that macrophages contribute to bone marrow tolerance in recurrent MI.

**Figure 8.**
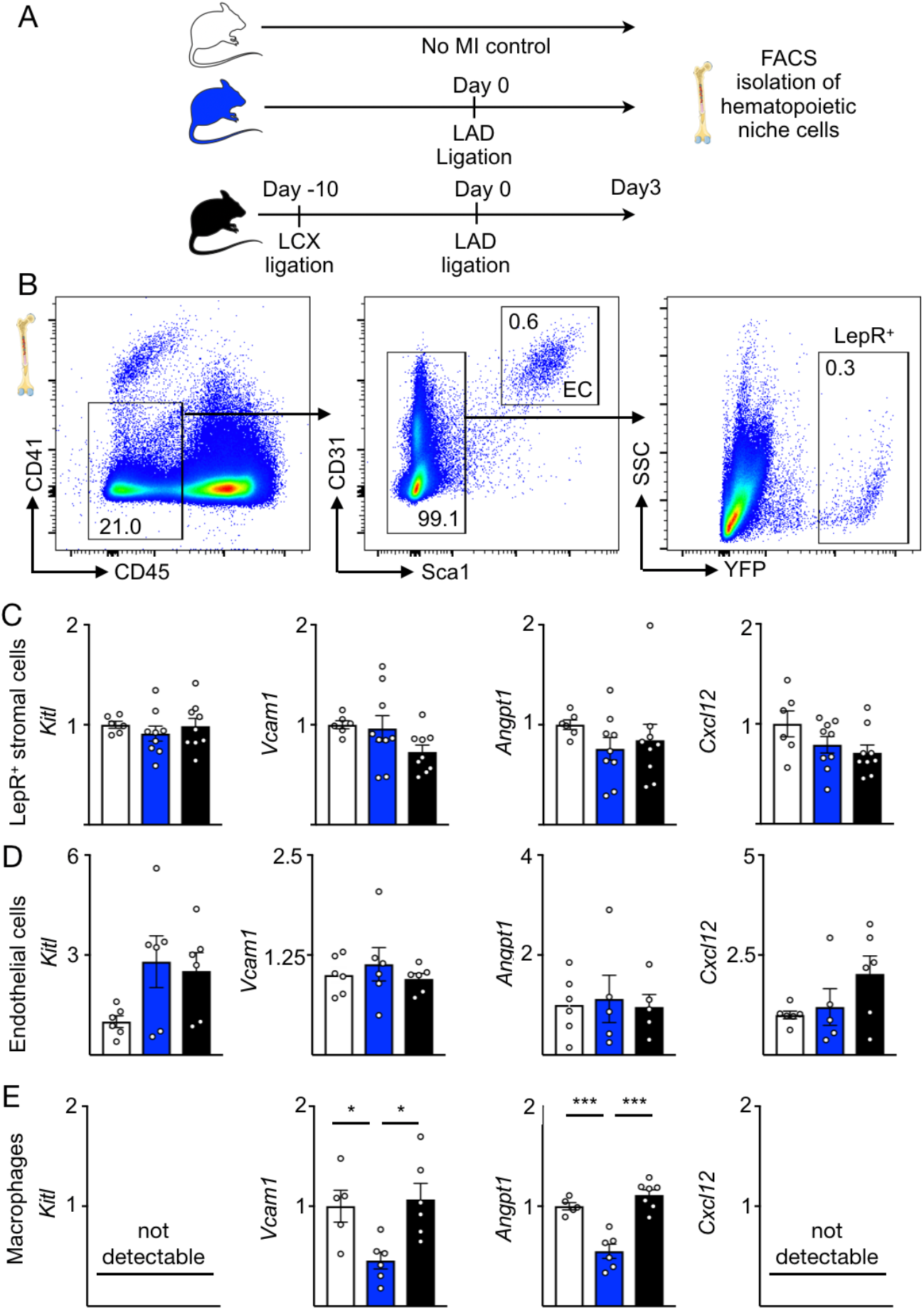
Macrophages attenuate myelopoiesis in recurrent MI by higher expression of quiescence- and retention-promoting factors. **(A)** Experimental design. Hematopoietic niche cells were isolated from bone marrow of naive mice and mice with either first or recurrent MI and subjected to gene expression analysis with qPCR. **(B)** Gating strategy for isolating bone marrow endothelial cells and LepR^+^ stromal cells. **(C-E)** Gene expression by qPCR in LepR^+^ stromal cells, endothelial cells and bone marrow macrophages. mRNA levels were normalized to *gapdh* Ct values. *p<0.05, **p<0.01, ***p<0.001, n=5-9, one-way analysis of variance (ANOVA) followed by Tukey’s multiple comparisons test. Data are mean ± s.e.m..

## Discussion

Innate immunity and cardiac inflammation have recently been linked to heart failure (Sager et al., 2016; Bajpai et al., 2018; Dick et al., 2019), a connection that may contribute to the dismal survival rate of patients with recurrent MI (Thune et al., 2011). There were no available systems for modeling inflammatory circuits in recurrent MI. We therefore designed a mouse model that allows the induction of two sequential infarctions in different anatomical locations of the same heart. The consequences were characterized by whole-heart fluorescent coronary angiography enabled by optical clearing of the organ and by cardiac MRI. Studying recurrent MI’s impact on innate immunity and emergency hematopoiesis, we observed a muted inflammatory response, which is induced by a circulating signal released after the first infarct and possibly mediated by bone marrow macrophages in the hematopoietic niche.

The heart recruits millions of circulating myeloid cells to the ischemic myocardium (Nahrendorf et al., 2007; Leuschner et al., 2012). Because these cells are short lived, with lifespans of about a day (Yona et al., 2013; Patel et al., 2017), the bone marrow augments leukocyte production to meet the organism’s need after MI in mice and humans (Dutta et al., 2012; Dutta et al., 2015b). After MI, the marrow is alerted by signals from the sympathetic nervous system (Dutta et al., 2012) and circulating mediators produced by the ischemic heart such as IL-1β and GM-CSF (Sager et al., 2015; Anzai et al., 2017). These mechanisms are not mutually exclusive and can either impact hematopoietic stem cells directly or relay their signals through bone marrow niche cells (Sager et al., 2015). This process controls HSPC quiescence, proliferation and differentiation, and subsequently blood leukocyte levels, in response to myocardial injury. Bone marrow activation is important not only in the context of acute MI. Indeed, chronic inflammatory processes in the cardiovascular system also rely on constant leukocyte supply from the bone marrow, which is stimulated by cardiovascular risk factors such as hyperlipidemia, chronic stress, insufficient sleep, hypertension and diabetes (Swirski et al., 2007; Yvan-Charvet et al., 2010; Nagareddy et al., 2013; Heidt et al., 2014; Al-Sharea et al., 2019; McAlpine et al., 2019).

Cardiovascular mortality correlates with blood leukocyte levels (Ernst et al., 1987; Maekawa et al., 2002; Madjid et al., 2004; Engstrom et al., 2009). As patients with recurrent MI have lower survival rates and aggravated heart failure (Thune et al., 2011), we speculated that recurrent MI would solicit heightened systemic supply of myeloid cells maintained by higher HSPC proliferation rates in the bone marrow. Yet our data indicate that in mice with recurrent MI the opposite is true. Peripheral blood monocytosis and myeloid cell numbers in the heart were lower, and bone marrow activity dimmed after recurrent MI. Clinical data clearly demonstrate that patients with recurrent MI have a worse prognosis than patients with only one infarct(Thune et al., 2011; Stone et al., 2014); however, information about blood leukocytes, infarct inflammation and bone marrow activity in this cohort are currently unavailable. This should be investigated to compare our findings in mice to human patients.

The potential clinical implications of this work are manifold: if recurrent MI leads to a muted and even insufficient inflammatory response, then perhaps these patients should not receive anti-inflammatory therapeutics after MI. The potential of such therapeutic interventions is currently under active discussion, especially in light of the recent CANTOS trial (Ridker et al., 2017). Prior studies exploring anti-inflammatory therapeutics neutralizing TNFα (Chung et al., 2003; Mann et al., 2004) or nonspecific drugs such as colchicine (Deftereos et al., 2014) returned discouraging results. Careful patient selection, including criteria such as first rather than recurrent MI, may help define a patient subpopulation that benefits from therapeutically dampened hematopoiesis. Determining the optimal inflammatory response required for proper infarct healing may explain how the phenotype we observed in mice links to patients with recurrent MI. If an infarct does not recruit a sufficient number of myeloid cells, wound healing is delayed, because dying cardiomyocytes will not be removed, resulting in more necrotic tissue remnants and lack of granulation tissue and collagen matrix (Nahrendorf et al., 2010). Patients with leukocytosis but also leukopenia have a higher mortality after acute MI (Grzybowski et al., 2004; Coller, 2005). Non-selectively depleting circulating monocytes and inhibiting myeloid cell recruitment to the heart impair wound healing and increase left ventricular remodeling after MI in wild type mice (Nahrendorf et al., 2007; van Amerongen et al., 2007). Thus, future studies should examine immune cell levels and hematopoiesis in patients with recurrent MI.

Regarding mechanisms that may attenuate bone marrow activation after recurrent MI, we identified altered expression of niche factors in bone marrow macrophages, which are known to influence HSPC retention, quiescence and differentiation (Chow et al., 2011; Chow et al., 2013; Hashimoto et al., 2013). In our experiments, recurrent MI did not result in a reduction of retention factors that is observed after a first infarct, which induces lower expression of Cxcl12, Vcam1, Scf and Angiopoietin1. Our parabiosis experiments indicate that a circulating, likely heart-derived factor is causally involved. Such signals, which have to be identified in future work, might act directly on macrophages in the bone marrow, where they could induce metabolic and epigenetic rewiring. In “trained immunity”, different stimuli induce epigenetic modifications involving histone modifications (Saeed et al., 2014) or metabolic changes that prime myeloid cells and their progenitors for secondary challenges (Cheng et al., 2014; Mitroulis et al., 2018), enabling a higher response after re-stimulation. Other pathways are involved in macrophage tolerance, which is mediated by LPS and requires TLR4 signaling (Foster et al., 2007; Netea et al., 2016). Therefore, it would be interesting to conduct epigenetic screens in bone marrow macrophages to compare the effects of first and recurrent MI. We expected to see an increased immune response in recurrent MI compared to a first MI, partially because the ischemic myocardium produces IL-1β and GM-CSF (Sager et al., 2015; Anzai et al., 2017), two cytokines that induce HSPC “training” rather than “tolerance” (Christ et al., 2018; Mitroulis et al., 2018). Surprisingly, our data show that MI induces “tolerance”. Potential candidates inducing such bone marrow tolerance after MI are the TLR4 ligands HMGB1 and S100, which are produced by ischemic myocytes and fibroblasts (Andrassy et al., 2008; Volz et al., 2012; Rohde et al., 2014).

In our studies described here, we examined young and otherwise healthy mice, which likely mount an optimal inflammatory response in response to myocardial injury (Nahrendorf et al., 2010). Increasing or lowering inflammation in these mice has negative effects on infarct healing. Patients with recurrent MI have atherosclerosis and often multiple cardiovascular risk factors, which all may give rise to low grade inflammation (Netea et al., 2017; Ridker et al., 2017). For instance, hyperlipidemia heightens the bone marrow’s inflammatory reactions to LPS (Christ et al., 2018). Accordingly, the effects of recurrent MI observed in healthy wild type mice may be different if atherosclerosis and its risk factors are present.

In summary, our data strongly suggest that MI alters the innate immune response, which affects systemic inflammation and the organism’s capacity to react to subsequent ischemic events and infections. The many preclinical questions arising from our observations can be tackled using the new mouse model of recurrent MI described in this Resource Article.

## Methods

### Animals

C57BL/6 (Jax No. 000664) and B6.SJL-*Ptprc*^*a*^ *Pepc*^*b*^/BoyJ (Jax No. 002014) were purchased from Jackson Laboratory. B6.129(Cg)-*Lepr*^*tm2(cre)Rck*^/J (Jax No. 008320) and B6.129X1-*Gt(ROSA)26Sor*^*tm1(EYFP)Cos*^/J (Jax No. 006148) were bred in house to generate stromal cell reporter mice. Genotyping for each strain was performed as described on the Jackson Laboratory website. 10-12-week old mice were used for experiments. All mice were maintained and bred in the pathogen-free environment of the Massachusetts General Hospital animal facility. Age-matched female mice were randomly allocated to either control or treatment groups. LPS (2μg, from Escherichia coli 055:B5, Sigma Aldrich) was administered intraperitoneally to mice that either had or had not previously received myocardial infarction surgery of the LAD. Clodronate and control liposomes were purchased from Liposoma Research. Depletion studies were performed by intraperitoneally injecting 100 μL of liposome formulation per mouse as recommended by the manufacturer.

### Procedures to induce myocardial ischemia

To permanently ligate the left anterior descending coronary artery (LAD), we followed previously published protocols(Eberli et al., 1998; Lutgens et al., 1999). Mice were anesthetized with 1.5-2% isoflurane supplemented with oxygen, intubated and ventilated (Inspira, Harvard Apparatus). After thoracotomy, the heart was exposed, and the left coronary artery was identified, and permanently ligated with a monofilament nylon 8-0 suture. The rib cage was then closed using two separate 5-0 sutures, with a 20G sheath left between the sutures. Finally, air was removed from the thoracic cavity via the sheath and a 3 ml syringe prior to extubation.

To permanently ligate the circumflex artery (LCX), the heart was exposed and the first artery branching off the LAD was located. If the LCX was not visible, which occurred in about half the cases, the ligation was conducted lateral of the LAD in close proximity to the left atrium. Successful induction of myocardial infarction was verified by identifying a pale ischemic area after ligation.

To induce recurrent MI, mice first received a LCX followed by a LAD ligation 10 days later. The chest was reopened and adhesions were carefully detached. The LAD was identified and permanently ligated, followed by chest cavity closure as described above. In mice subjected to ischemia-reperfusion injury, the coronary artery was ligated for 30 minutes, followed by ligature removal.

### Parabiosis

Parabiosis surgery was performed as previously described(Robbins et al., 2013). In brief, after shaving the corresponding lateral aspects of a CD45.1 and a CD45.2 mouse, matching skin incisions were made in each mouse from behind the ear to the tail, and the subcutaneous fascia were bluntly dissected to create 0.5 cm of free skin. The left olecranon of one mouse was sutured to the right olecranon of the other mouse with 3.0 non-absorbable suture. Afterwards, the left knee joint of the first mouse was connected to the right knee joint of the second mouse in the same way. Finally, the dorsal and ventral skins were approximated by continuous suture with a 5.0 absorbable Vicryl suture. Mice were joined for two weeks. The CD45.2 mouse was then subjected to MI. After five days, parabionts were surgically split and the CD45.1 mouse received MI surgery five days later.

### Optical clearing and fluorescence coronary angiography (FCA)

For in vivo vasculature labeling, mice were anesthetized with 1.5-2% isoflurane supplemented with oxygen and sacrificed by perfusion through the left ventricle with 20 mL of ice-cold PBS followed by 20 mL of 4% formaldehyde solution in PBS (Thermo Scientific, Waltham, MA, USA). A 2% (w/v) solution (10 mL) of porcine skin gelatin was prepared in boiling PBS. After the solution reached a temperature below 40°C, it was combined with a 0.5% (w/v) fluorescein-albumin, filtered and injected in the mouse left ventricle(Tsai et al., 2009). After perfusion the heart was resected and placed in an ice-cold PBS solution for one hour. The heart was then fixed for one hour in a 4% formaldehyde solution, before being washed for 30 min in PBS and imaged. Whole heart optical clearing was then performed using the CUBIC (clear, unobstructed brain imaging cocktails and computational analysis) method(Susaki et al., 2014). Hearts were immersed for one day in a chemical mixture obtained by mixing 25 wt% urea (U16–3, Fisher Scientific Hampton, NH, USA), 25 wt% N,N,N0,N0-tetrakis(2-hydroxypropyl) ethylenediamine (50-014-48142, Fisher Scientific) and 15 wt% Triton X-100 (85111, Life Technologies, Carlsbad, CA, USA) at room temperature. Hearts were imaged using a 2x objective with a confocal microscope (FV1000, Olympus America). To visualize the vasculature network along the curvature of the heart, images were displayed as maximum intensity projection of z-stack datasets.

### MRI

MRI was carried out in naive mice and after permanent LCX and LAD ligations at the indicated time points (see Figure 1G) as previously described(Panizzi et al., 2010; Sager et al., 2015). We obtained cine images of the left ventricular short axis by using a 4.7 Tesla horizontal bore Pharmascan (Bruker) and a custom-built mouse cardiac coil (Rapid Biomedical) and a cine gradient echo FLASH-sequence with IntraGate, a self gating software to compensate respiratory and cardiac motion. Acquisition parameters were: echo time: 2.94 ms, repetition time 12 ms, matrix size 200 x 200 x 1, voxel size 0.15 mm x 0.15 mm x 1 mm, oversampling 350 and 16 cine frames. Delayed enhancement imaging was done 10-20 minutes after i.v. injection of 0.3 mmol/kg Gd-DTPA at the day 3 timepoint. Images were analyzed using the software Horos (http:// https://horosproject.org). Left ventricular volume, ejection fraction and infarct size were acquired and calculated as previously(Panizzi et al., 2010; Sager et al., 2015).

### TTC staining

Hearts were harvested and sectioned with a heart slicer (zivic instruments). Slices were then incubated in a solution of 2% TTC (2,3,5-Triphenyltetrazolium chloride, Sigma-Aldrich) for 15 minutes protected from light. ImageJ software was used to quantitate infarct size (imagej.nih.gov).

### Tissue processing

Peripheral blood was collected by retro-orbital bleeding using heparinized capillary tubes (BD Biosciences). For flow cytometry analysis, red blood cells were lysed with 1x red blood cell lysis buffer (BioLegend). For organ harvest, mice were perfused through the left ventricle with 20 mL of ice-cold PBS. After harvest, bone marrow for qPCR analysis was isolated by centrifugation. The metaphysis of one end of the tibia was removed, and the bone was spun at 6000g with this ‘open end’ facing down. For qPCR analysis, bone marrow was stored in RLT buffer (Qiagen) at −80°C for further analysis. For flow cytometry analysis of HSPC and mature leukocytes, bone marrow from the femurs was flushed with FACS buffer (1x PBS supplemented with 0.5% BSA). Infarct heart tissue was minced into small pieces and subjected to enzymatic digestion with 450 U/mL collagenase I, 125 U/mL collagenase XI, 60 U/mL DNase I and 60 U/mL hyaluronidase (all Sigma-Aldrich) for 30 minutes at 37°C under agitation (750rpm). Digested tissues were then triturated and filtered through a 40μm nylon mesh (Falcon), washed and centrifuged to obtain single-cell suspensions. For stromal cell isolations, the bone marrow fraction was digested to isolate endothelial cells and LepR^+^ cells. The bone marrow fraction was isolated from the long bones and pelvis of mice by flushing the bone marrow plug with a syringe into PBS supplemented with 2% FBS. After the bone marrow plug sank to the bottom of the tube, the supernatant was removed and replaced with the digestion mix: collagenase IV at 1mg/ml (Sigma Aldrich, C5138), Dispase at 2mg/ml (Gibco by Life technology, 17105-041) and DNAse I (Thermo scientific, 90083) in HBSS buffer (Gibco by Life Technology, 14025-092). The bone marrow plug was digested 3 x 15 minutes at 37°C. Bone stromal cells in suspension were washed with PBS supplemented with 2% FBS.

### Flow cytometry

All single cell suspensions were stained at 4°C in 300μl FACS buffer (1x PBS supplemented with 0.5% BSA). For HSPC staining, isolated bone marrow cells were first stained with biotin-conjugated anti-mouse antibodies directed against mouse hematopoietic lineage markers, including CD3 (100304), CD4 (100404), CD8 (100704), CD49b (108904), CD90.2 (105304), CD19 (115503), B220 (103204), NK1.1 (108704), TER119 (116204), CD11b (101204), CD11c (117304), Gr1 (108404; all 1:300, BioLegend). This was followed by a second staining with antibodies for CD16/32-BV711 (101337), CD34-FITC (553733, BD Biosciences), CD135-PE (135306), CD48-AF700 (103426), CD115-BV421 (135513), CD150-PerCP/Cy5.5 (115922), c-kit-PE/Cy7 (105814), Sca-1-BV605 (108133), streptavidin-APC/Cy7 (405208; all 1:150, BioLegend unless otherwise indicated) and, where applicable, BrdU-APC (1:50, BD Biosciences).

For cardiac leukocyte staining, cells were stained with PE-conjugated mouse hematopoietic lineage markers, including B220 (103208), CD103 (121406), CD19 (115508), CD3 (100206), CD49b (108908), CD90.2 (140308), NK1.1 (108708) and Ter119 (116208; all 1:300, BioLegend) as well as CD11b-APC (1:300, 101212), CD45-BV711 (1:300, 103147), F4/80-PECy7 (1:150, 123114), Ly-6C-BV605 (1:300, 128035) and Ly-6G-FITC (1:300, 127605, all BioLegend). To stain mature leukocytes in the bone marrow, cells were stained with B220-APC/Cy7 (1:300, 103223), CD3-APC/Cy7 (1:300, 100221), Gr1-PE (1:300, 108407), CD115-BV605 (1:300, 135517) and F4/80 APC (1:150, Cl A3-1, AbD Serotec). Isotype controls for the APC antibody were used to set the gate for F4/80^pos^ macrophages. To profile macrophages in the bone marrow, we used antibodies for VCAM1-PerCP/Cy5.5 (1:150, 105715), CX3CR1-PerCPCy5.5 (1:300,149009), I-A/I-E-PacificBlue (1:150, 107619) and CD169-PeCy7(1:150, 142411). Respective isotype controls were used. All BioLegend unless otherwise indicated. For blood leukocyte staining, cells were stained with CD19-BV711 (11555), CD45.2-APC-Cy7 (109803) or CD45.1-APC-Cy7 (110715), CD90.2-PE (140308), Ly-6C-BV605 (128035), Ly-6G-FITC (127605), CD11b-APC (101212), CD115-BV421 (135513) and CX3CR1-PerCP (149009, all 1:300, BioLegend). Blood and bone marrow leukocyte staining samples were fixed with BD Cytofix (BD Biosciences) and analyzed within 24 hrs. BrdU staining (552598, BD Biosciences) was done according to the manufacturer’s protocol.

### Gating strategy for flow cytometry

All cells were pre-gated on single cells (FSC-A vs FSC-W, and SSC-A vs SSC-W). LSK were identified as Lin^−^ c-kit^+^ Sca-1^+^. These were further divided into long-term hematopoietic stem cells (LT-HSCs; Lin^−^ c-kit^+^ Sca-1^+^ CD150^+^ CD48^-^) and multipotent progenitors (MPPs; Lin^−^ c-kit^+^ Sca-1^+^, CD135^−^ CD150^−^ CD48^+^). Granulocyte macrophage progenitors (GMP) were identified as Lin^−^ c-kit^+^ Sca-1^−^ CD16/32^+^ CD34^+^. Marrow monocytes were identified as CD3^−^ B220^−^ Gr1^+^ CD115^+^, marrow neutrophils as CD3^−^ B220^−^, Gr1^+^ CD115^−^ and bone marrow macrophages as CD3^−^ B220^−^ Gr1^−^CD115^−^ F4/80^+^ SSC^lo^. Blood monocytes were identified as CD45^+^ CD19^−^ CD90.2^−^ Ly-6G^−^ CD11b^+^ CD115^+^ and neutrophils as CD19^−^ CD90.2^−^ CD115^−^ CD11b^+^ Ly-6G^+^. Leptin receptor (LepR^+^) cells were identified as CD41^−^ CD45^−^ Ter119^−^ Sca1^−^ CD31^−^ YFP^+^ in the bone marrow of LeptinRcre-EYFP mice. Endothelial cells were identified as CD41^−^ CD45^−^ Ter119^−^ CD31^+^ Sca1^+^ in the bone marrow of LepR-YFP mice. For compensation, the aforementioned antibodies were conjugated to OneComp eBeads (Affymetrix Inc). Unstained and YFP^+^ control samples were used for compensation and to control the gating strategy. All data were acquired on an LSRII (BD Biosciences) and analyzed with FlowJo software.

### Cell sorting

To purify bone marrow stromal cells, samples were stained with CD31-PE (102507), CD41-PerCP (133917), CD45.2-BV605 (103116), Ter119-APC/Cy7 (116223) and Sca1-PeCy7 (108113, all 1:100, Biolegend) and FAC-sorted with a FACSAria II cell sorter (BD Biosciences). For bone marrow macrophages, the same staining panel was used as for bone marrow leukocytes. All cells were pre-gated on single cells (as determined by FSC-A vs FSC-W, and SSC-A vs SSC-W) and viable cells (FxCycle™ Violet Stain^−^).

### Real-time qPCR

Total RNA from bone marrow was extracted using the RNeasy Mini kit (Qiagen) according to the manufacturer’s protocol. For bone marrow stromal cells, the NucleoSpin RNA XS kit (Takara Bio) was used. RNA (1μg) was reverse-transcribed using the High-Capacity RNA-to-cDNA kit (Qiagen) according to the manufacturer’s instructions. For stromal cells, all isolated RNA was transcribed. TaqMan gene expression assays were used to quantify target genes. The relative changes were normalized to *Gapdh* mRNA levels using the 2^−ΔΔCT^ method.

### Enzyme-linked immunosorbent assay

For troponin-I ELISA (Life diagnostics, CTNI-1-US) measurements, mice were bled retroorbitally 24h after induction of myocardial infarction. Blood plasma was obtained by centrifugation. Plasma was 4x diluted. Samples were analyzed according to the manufacturers’ instructions.

### Statistical analyses

Statistical analyses were conducted with GraphPad Prism software Version 8.0. Results are depicted as mean ± standard error of mean. Data were first tested for normality by D’Agostino-Pearson omnibus normality test. For a two-group comparison, a Student’s t-test was applied if the pretest for normality (D’Agostino-Pearson normality test) was not rejected at the 0.05 significance level; otherwise, a Mann-Whitney U test for nonparametric data was used. To compare more than two groups, an ANOVA test, followed by a Tukey test for multiple comparison, was applied. *P* values of <0.05 indicate statistical significance. No statistical method was used to predetermine sample size and animals were randomly assigned to treatment groups.

## Acknowledgements

This work was funded in part by the National Institutes of Health HL139598, HL125428, HL131495, HL131478, T32HL076136, the European Union’s Horizon 2020 Research and Innovation Program under grant agreement No 667837 and the MGH Research Scholar Program. S. Cremer and D. Rohde were funded by Deutsche Forschungsgemeinschaft (CR603/1-1 and RO5071/1-1). The authors declare no competing financial interests. We acknowledge Kaley Joyes for editing the manuscript.

## Author contributions

S. Cremer and M.J. Schloss developed and validated the mouse model of recurrent MI. C. Vinegoni developed fluorescent coronary angiography. S. Cremer, M.J. Schloss, S. Zhang, D. Rohde and C.Vinegoni performed experiments and collected, analyzed and discussed data. G. Wojtkiewicz and S. Schmitt performed imaging experiments and collected data. S. Cremer, F. Swirski and M. Nahrendorf conceived experiments and discussed results and strategy. M. Nahrendorf conceived, designed and directed the study. S. Cremer and M. Nahrendorf wrote the manuscript, which was revised and approved by all authors.

---

**Supplementary Figure 1.**
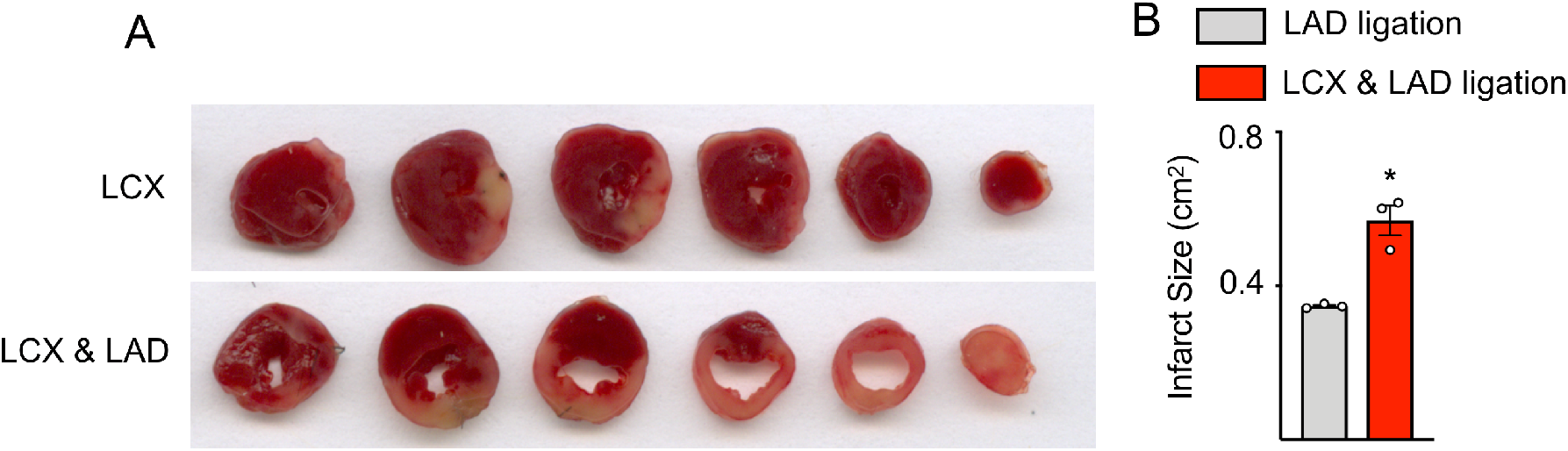
2,3,5-Triphenyltetrazolium chloride (TTC) staining. **(A)** TTC staining of a heart with an isolated LCX ligation and both ligations. **(B)** Quantification of infarct size assessed by TTC staining of LAD MIs compared to hearts in which both vessels were ligated. *p<0.05, n=3, Student’s t-test. Data are mean ± s.e.m..

**Supplementary Figure 2.**
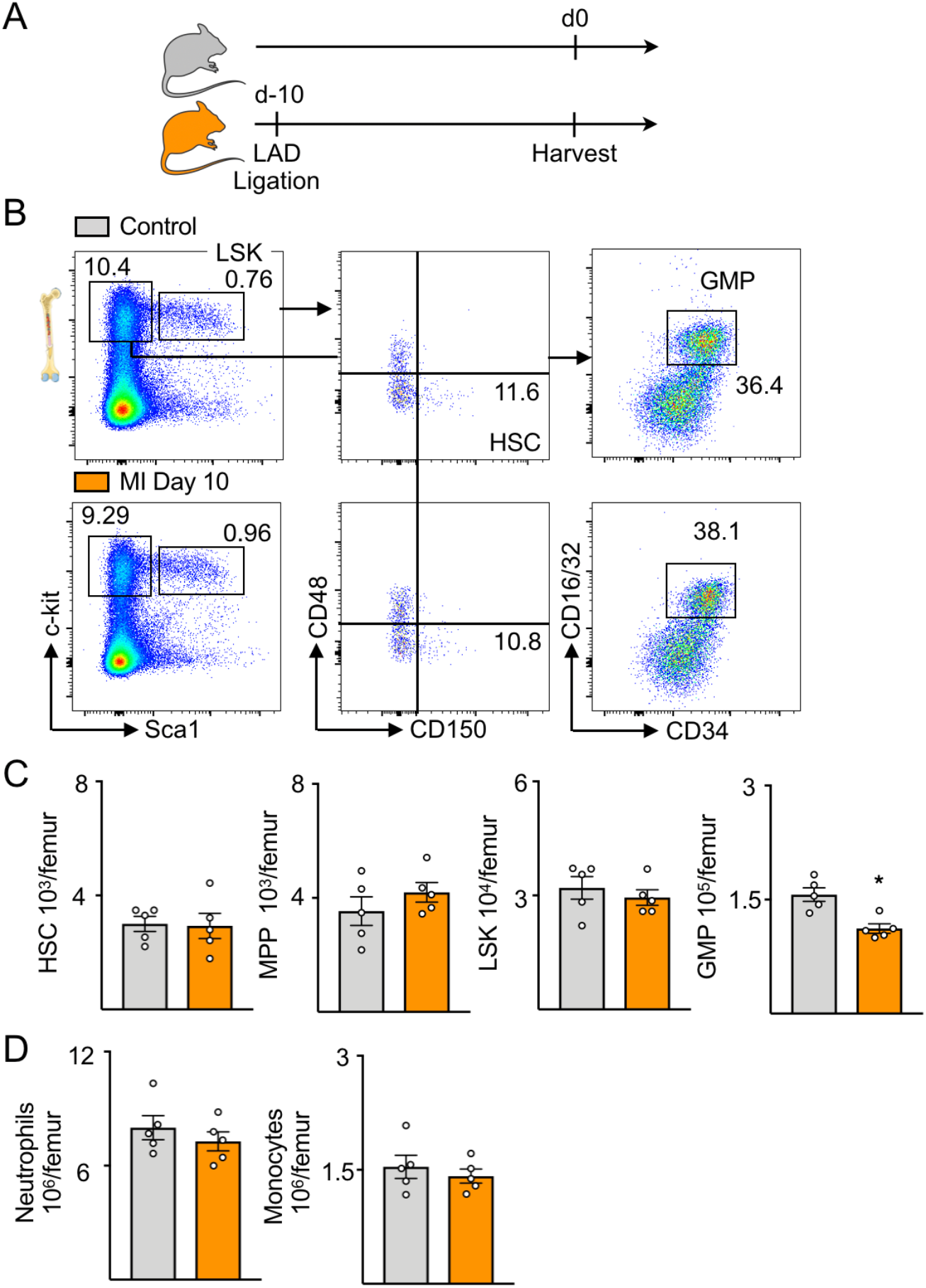
HSPC numbers 10 days after MI. **(A)** Experimental outline. Mice had LAD MI 10 days before bone marrow analysis. **(B)** Dot plots and **(C)** quantification of HSPC in bone marrow of mice with MI compared to naive mice. **(D)** Numbers of mature leukocytes in bone marrow. *p<0.05, n=5 Student’s t-test. Data are mean ± s.e.m..

**Supplementary Figure 3.**
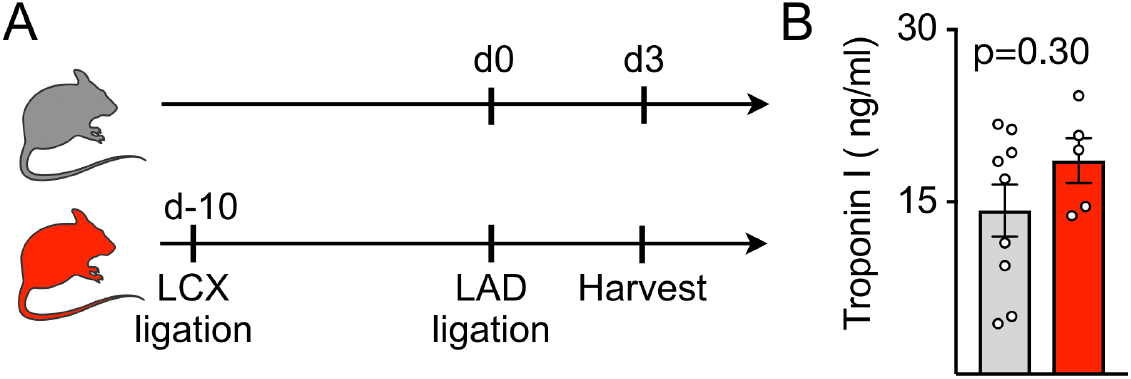
Troponin after a first versus recurrent MI. **(A)** Experimental design. **(B)** Troponin levels after LAD ligation as the first compared to recurrent MI. n=5-9.

**Supplementary Figure 4.**
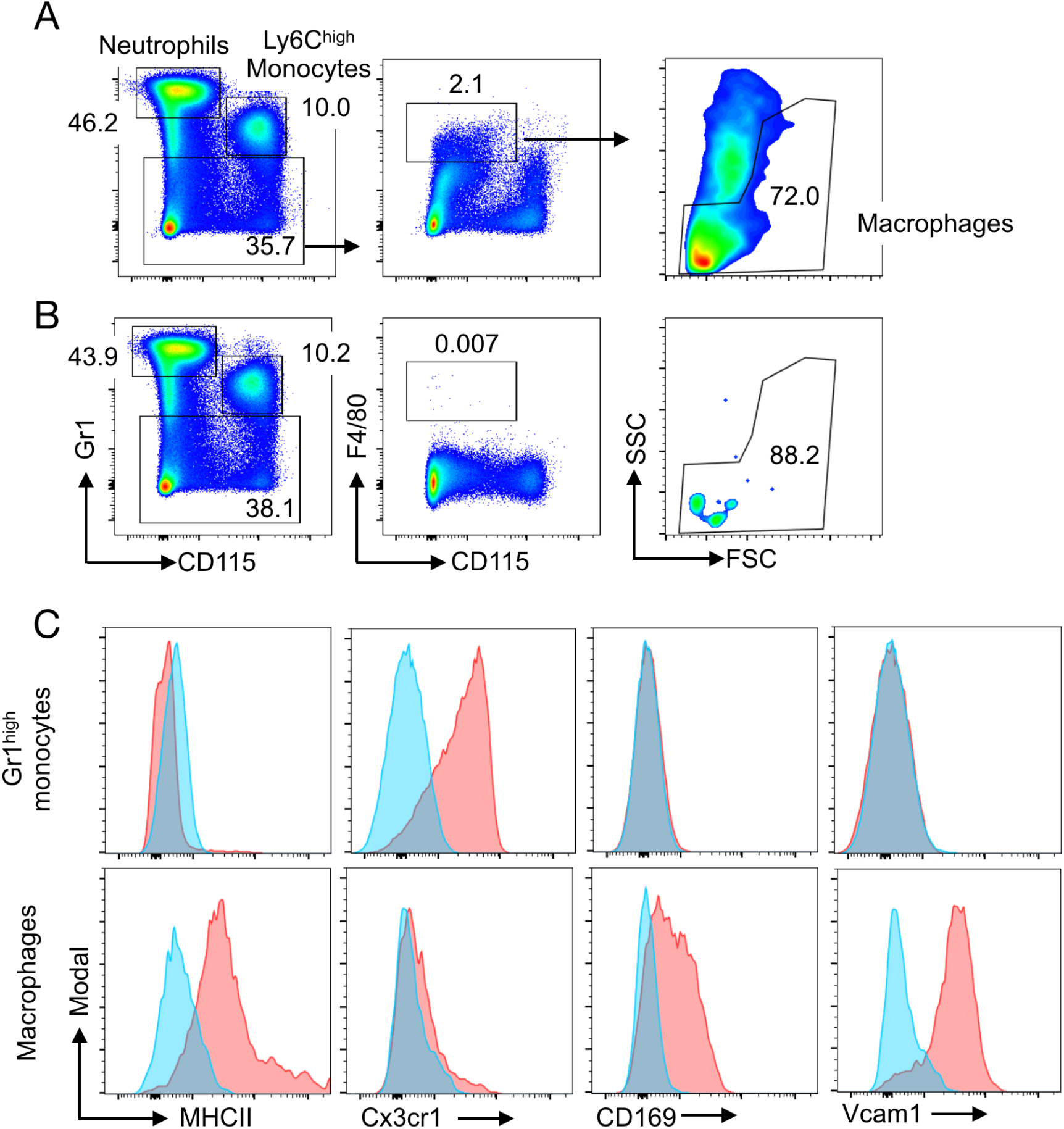
Macrophage gating. **(A)** Gating strategy for bone marrow macrophages with antibody directed against F4/80 and **(B)** isotype control. **(C)** Histograms in red depicting the expression of MHCII, Cx3cr1, CD169 and Vcam1 in Gr1^hi^ monocytes and bone marrow macrophages. Isotype controls are shown in blue.

**Supplementary Video 1: MRI short axis cine loop of LCX infarct.** Delayed enhancement MRI short axis views of a mouse with LCX MI demonstrating injury of posterolateral left ventricular wall.

**Supplementary Video 2: MRI short axis cine loop after recurrent MI (LCX followed by LAD ligation).** Delayed enhancement MRI short axis views of the same mouse as in supplementary video 1, now also showing injury of the anterior left ventricular wall.

